# Harnessing DSB repair to promote efficient homology-dependent and -independent prime editing

**DOI:** 10.1101/2021.08.10.455572

**Authors:** Martin Peterka, Nina Akrap, Songyuan Li, Sandra Wimberger, Pei-Pei Hsieh, Dmitrii Degtev, Burcu Bestas, Jack Barr, Stijn van de Plassche, Patricia Mendoza-Garcia, Saša Šviković, Grzegorz Sienski, Mike Firth, Marcello Maresca

**Affiliations:** Genome Engineering, Discovery Sciences, BioPharmaceuticals R&D Unit, AstraZeneca, Gothenburg, Sweden; Department of Chemistry & Molecular Biology, University of Gothenburg, Gothenburg, Sweden; Data Sciences and Quantitative Biology, Discovery Sciences, AstraZeneca, Cambridge, UK

## Abstract

Prime editing recently emerged as a next-generation approach for precise genome editing. Here we exploit DNA double-strand break (DSB) repair to develop two novel strategies that install precise genomic insertions using an *Sp*Cas9 nuclease-based prime editor (PEn). We first demonstrate that PEn coupled to a regular prime editing guide RNA (pegRNA) efficiently promotes short genomic insertions through a homology-dependent DSB repair mechanism. While PEn editing lead to increased levels of by-products, it rescued pegRNAs that performed poorly with a nickase-based prime editor. We also present a small molecule approach that yielded increased product purity of PEn editing. Next, we developed a homology-independent PEn editing strategy by engineering a <u>s</u>ingle <u>pr</u>imed <u>in</u>sertion <u>gRNA</u> (springRNA) which installs genomic insertions at DSBs through the non-homologous end joining pathway (NHEJ). Lastly, we show that PEn-mediated insertions at DSBs prevent Cas9-induced large chromosomal deletions and provide evidence that continuous Cas9-mediated cutting is one of the mechanisms by which Cas9-induced large deletions arise. Altogether, this work expands the current prime editing toolbox by leveraging distinct DNA repair mechanisms including NHEJ, which represents the primary pathway of DSB repair in mammalian cells.

## Introduction

Proper utilization of cellular DNA repair mechanisms is instrumental to any successful genome editing strategy. The activity of different DNA repair pathways is highly dependent on tissue and cell type, chromatin context, and the DNA sequence of the target locus ^1–4^.

Targeted DNA insertions represent a particularly challenging type of precise genome modification but have the potential to repair the majority of genetic diseases. A common approach to introduce DNA insertions is to induce a targeted DNA double-strand break (DSB) using a site-specific nuclease combined with the delivery of a donor DNA repair template to stimulate homology-directed repair (HDR) at the targeted locus. A major disadvantage of this strategy is the limited activity of homologous recombination, which is restricted to S/G2 phases of the cell cycle and is generally absent in postmitotic cells ^5^.

Unlike homologous recombination, DNA end joining repair mechanisms such as non-homologous end joining (NHEJ) or alternative end joining (a-EJ) pathways remain active throughout the cell cycle and act as the major pathways of DSB repair in mammalian cells ^6–8^. While being typically considered error prone, NHEJ can repair DSBs with high fidelity ^9,10^. In contrast, the homology-dependent a-EJ pathway leads to deletions, which are highly predictable ^11,12^. The respective precision and predictability of NHEJ and a-EJ have been successfully exploited for precise genome modifications including DNA insertions ^13–16^. Harnessing DNA end joining pathways represents a valuable genome editing strategy, because most adult tissues are comprised of postmitotic cells unable to perform homologous recombination ^3,17^.

The recently developed CRISPR-based prime editing can install a wide spectrum of genomic modifications including deletions, substitutions and insertions without the need of a separate DNA template and without introducing DSBs ^18^, therefore offering a major advantage over existing genome editing methods. The PE2 prime editor combines Cas9 (H840A) nickase with an engineered reverse transcriptase (RT) to install an edit encoded directly in the <u>p</u>rime <u>e</u>diting <u>g</u>RNA (pegRNA). The cascade of events leading to a successful prime editing outcome is comprised of 1) Cas9-mediated nicking of the target site 2) hybridization of the pegRNA-encoded primer binding site (PBS) to the 3’ end of the nick 3) pegRNA-templated extension of the primed 3’ end of the nick by RT resulting in a “flap” containing the desired edit, and 4) hybridization and ligation of the flap with the targeted locus. DNA repair mechanisms responsible for the successful incorporation of the 3’ flap remain to be described in detail and might not be universally available in different cellular and genomic contexts, potentially limiting the scope and efficiency of the nickase-based PEs. A recent report suggests a possible dependency of PE2 editing on cell cycle progression reminiscent of HDR ^19^. Thus, a prime editing strategy harnessing a wider spectrum of DNA repair pathways would be a valuable addition to the prime editing toolbox.

Here we introduce a new prime editor, Prime Editor nuclease (PEn), that combines RT and the wild-type *Sp*Cas9 nuclease and show that prime editing can be performed at DSBs by utilizing DNA end joining repair pathways. We present two PEn strategies to robustly install small insertions via distinct DNA repair mechanisms. The first strategy utilizes regular pegRNAs to promote small DNA insertions by a homology-dependent DSB repair mechanism. This strategy worked robustly across different genomic loci as well as with pegRNAs displaying inefficient editing when combined with PE2. The second strategy relies on a novel sgRNA design to install small insertions through precise NHEJ. We also present a small molecule approach to decrease unintended by-products of PEn editing. Finally, we show that unlike editing with Cas9 alone, PEn does not induce large unintended on-target deletions, likely because PEn-mediated insertions at DSBs prevent NHEJ-mediated restoration of the wild-type sequence at the target locus. This suggests that the futile cycle of nuclease-mediated cut and NHEJ-mediated precise repair may be a possible cause of DSBs genotoxicity associated with Cas9 treatment.

## Results

### *SpCas9* nuclease-based prime editing

To test if a Cas9 nuclease-based prime editor can install small insertions at DSBs through a DNA end joining repair mechanism, we reverted the Cas9(H840A)-based PE2 into wtCas9-PE, designated here as PEn. We have constructed pegRNAs encoding small insertions of various sizes (6-18 bp) (Supplementary table 1) against 10 genomic target sites and co-transfected HEK293T cells with a pegRNA and either PEn or PE2. NGS analysis of the editing outcomes at the targeted sites revealed successful intended insertions with varying frequencies and product purities for both PEn and PE2 (Figure 1a). We classified the edited alleles into three categories; 1) all prime edits, representing any type of RT-templated insertions 2) precise prime edits, that represent RT-templated insertions of intended size and 3) other indels. As expected, PE2 editing resulted in high product purity, but also showed large site-to-site variability of insertion efficiency. PEn editing resulted in variable rates of precise prime edits but in general higher as compared to PE2. Interestingly, PEn installed insertions efficiently even with pegRNAs that were suboptimal for PE2 (Figure 1a, *CTLA4*, *PDCD1*). We observed a similar trend in HeLa and HCT116 cells, where we tested PE2-optimal (*AAVS1*) as well as suboptimal (*CTLA4*) pegRNAs (Supplementary Figure 1). As expected, due to the use of wild-type Cas9 that cuts both DNA strands, we also observed variable levels of PEn-induced imprecise prime edits and indels. Alignments of PEn-edited reads revealed that most imprecise prime edits represent additional integrations matching RT templates (Figure 1b).

**Figure 1.**
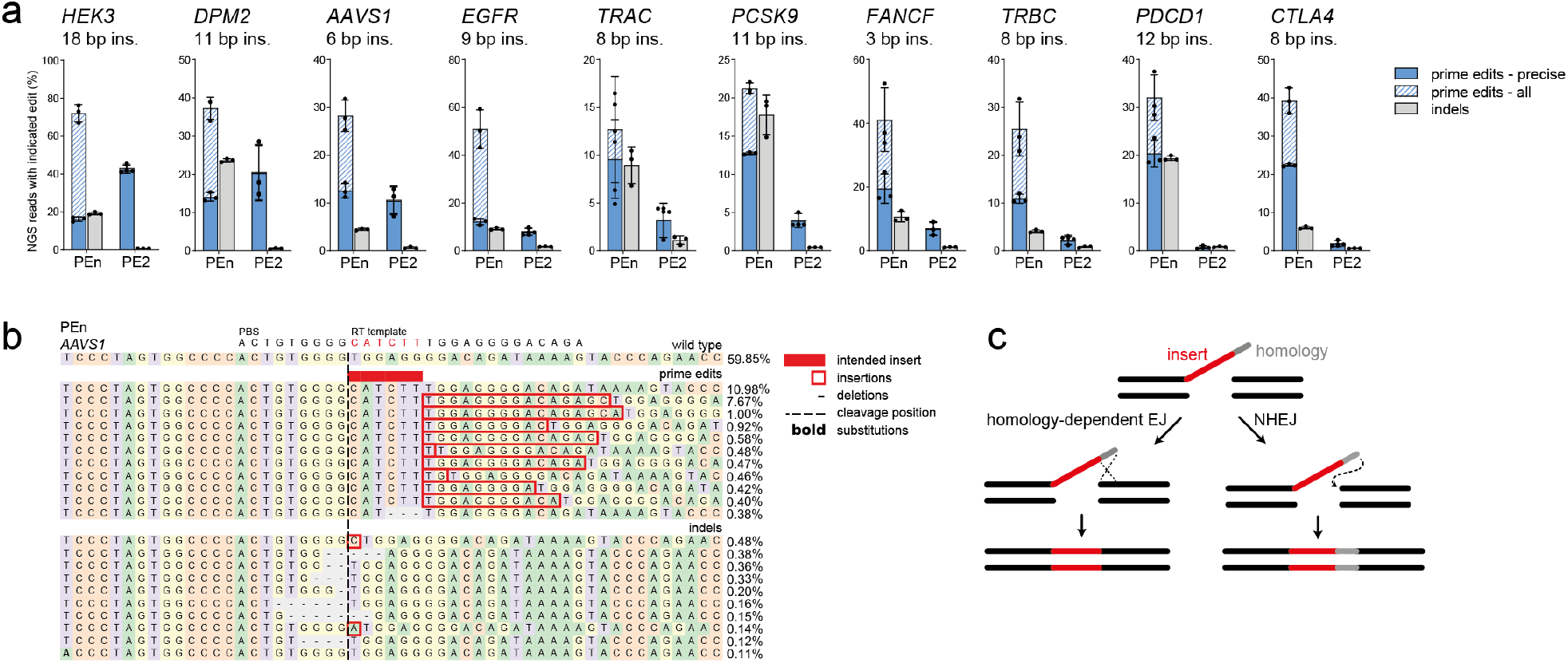
*Sp*Cas9 nuclease-based prime editing. a) NGS analysis of PEn or PE2-mediated targeted DNA insertions of indicated sizes using 10 different pegRNAs targeting endogenous loci in HEK293T cells. Plots show mean ± SD of 3 independent biological replicates. “prime edits – all” and “prime edits – precise” categories are superimposed. b) Representative alignment and allele frequencies of *AAVS1* edited with PEn and the indicated RT template in HEK293T cells. For each category, the top 10 variants are shown with a minimum frequency of 0.1%. c) Model of homology-dependent and NHEJ modes of PEn-mediated insertions at DSBs. Only DNA intermediates are shown.

### Mechanism of PEn-based prime editing

We reasoned that the integrated RT-templated homology tails might be products of DSB repair mediated by non-homologous end joining (NHEJ), while the precise insertions could occur through a homology-dependent process (Figure 1c). If true, the inhibition of NHEJ could shift the outcomes of PEn editing by decreasing the frequency of imprecise prime edits. To test this, we performed PEn editing in HEK293T cells treated with AZD7648, a small molecule inhibitor of DNA-PK, an essential mediator of NHEJ ^20^. Indeed, upon DNA-PK inhibition, the additional RT template integrations were abolished, and the remaining prime edits represented almost exclusively insertions of intended sizes (Figure 2a, b). At several loci, DNA-PK inhibitor treatment also led to an increase of total rates of correct insertions (Figure 2a – *DPM2*, *AAVS1*, *EGFR*).

**Figure 2.**
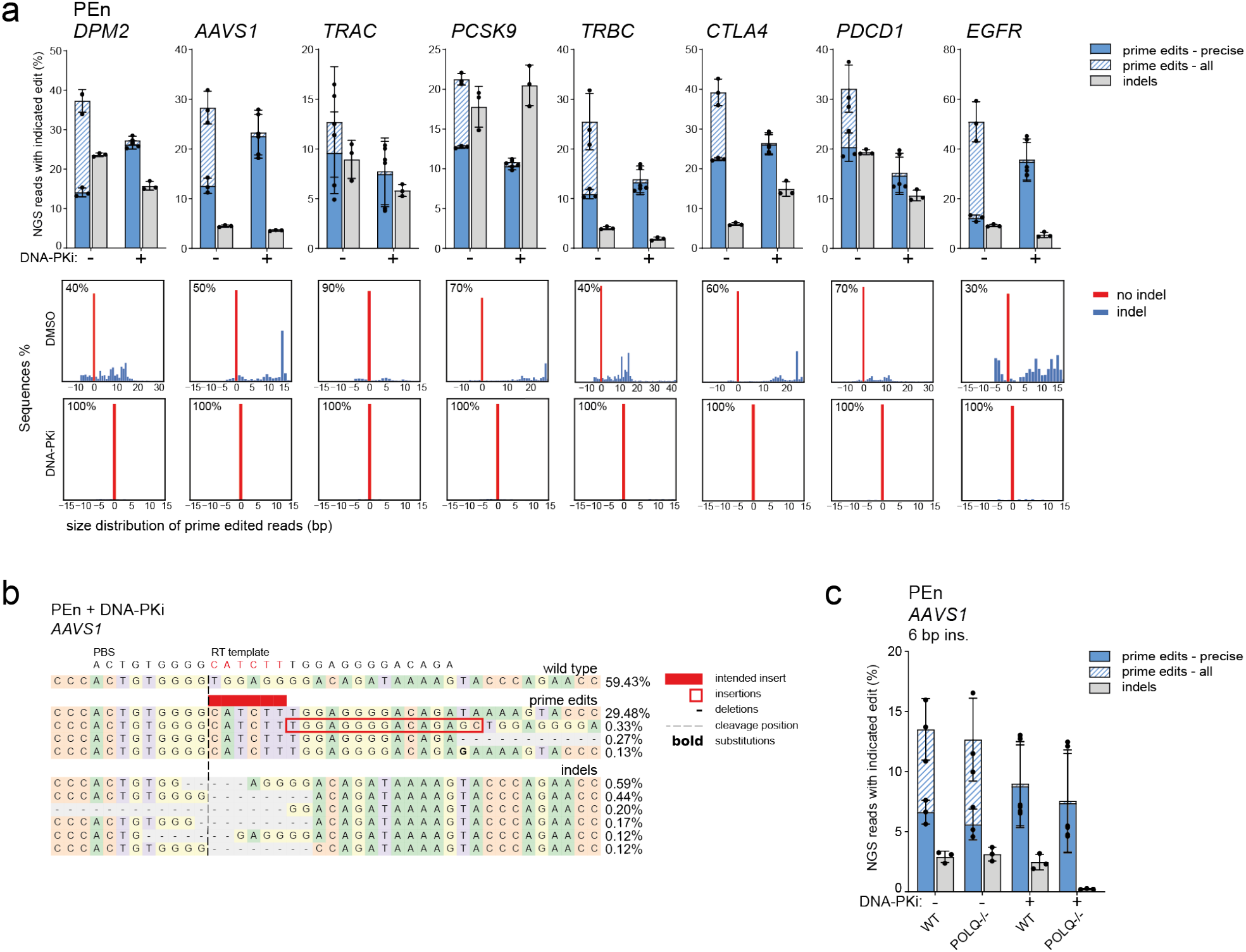
Mechanism of PEn-based prime editing. a) PEn editing using 8 different pegRNAs targeting endogenous loci in HEK293T cells treated with DNA-PK inhibitor or DMSO. Plots show mean ± SD of 3 independent biological replicates. “prime edits – all” and “prime edits – precise” categories are superimposed. Histograms below the bar plots represent percentages and size distribution of prime edited alleles for each target site with or without DNA-PK inhibition. b) Representative alignment and allele frequencies of the *AAVS1* locus edited with PEn and the indicated RT template in HEK293T cells treated with DNA-PK inhibitor. For each category, the top 10 variants are shown with a minimum frequency of 0.1%. c) NGS analysis of PEn editing outcomes of *AAVS1* in wild-type and *POLQ−/−* HEK293T cells with or without DNA-PK inhibitor treatment. The plot shows mean ± SD of 3 independent biological replicates.

Having pinpointed the contribution of NHEJ to PEn editing, we then investigated the mechanisms responsible for the homology-dependent DSB repair resulting in precise insertions. We reasoned that the short homology tails used in our pegRNA designs could utilize the a-EJ pathway, which typically uses DSB-proximal homology regions approximately 2-20 bp in length ^7^. To test this hypothesis, we performed PEn editing using the AAVS1 pegRNA with a 13 nt homology tail in HEK293T cells deficient in DNA Polymerase θ (encoded by *POLQ* gene), a crucial mediator of a-EJ ^7^. First, to confirm a-EJ inhibition in *POLQ*−/− background, we performed Cas9 editing of the *AAVS1* locus in these cells with or without DNA-PK inhibition. As expected, no indels were detected at the targeted site upon DNA-PKi treatment of *POLQ*−/− cells (Supplementary Figure 2), suggesting both NHEJ and e-EJ pathways were disabled. Despite this, PEn-mediated editing still proceeded efficiently, suggesting a mechanism independent of a-EJ (Figure 2c).

Altogether, our data reveal that the imprecise PEn prime edits are mediated by NHEJ. Accordingly, DNA-PK inhibition improves the purity of PEn editing and leads to increased efficiency in a locus-dependent fashion. Interestingly, PEn-mediated precise insertions appear to be independent of the Polymerase θ-mediated a-EJ pathway.

### PEn editing through NHEJ

The fact that PEn can install insertions via NHEJ, prompted us to test whether a pegRNA design encoding the intended insertion but no homology tail could still perform precise primed insertions through NHEJ-mediated integration (Figure 3a). This strategy could only be exploited to promote insertions at the cleavage site due to the lack of the homology region in the RT template. To test this <u>pr</u>imed <u>ins</u>ertions strategy (PRINS), we removed the homology regions from the RT templates of AAVS1 and CTLA4 pegRNAs, resulting in gRNAs with an extension containing only PBS and an intended insertion (<u>s</u>ingle <u>pr</u>imed <u>in</u>sertion <u>gRNA</u>, springRNA). PRINS editing with AAVS1 springRNA was able to install intended insertions in HEK293T, HeLa and HCT116 cells with up to 50% efficiency (Figure 3b). PRINS editing was completely abrogated by DNA-PK inhibition, confirming that NHEJ is responsible for the primed integration, (Figure 3b). The imprecise insertions constituted either truncated inserts or inserts longer than the intended size due to integrations of the gRNA scaffold sequence of various lengths (Figure 3c). The unintended edits were more frequent at *CTLA4*, where the majority of inserts were either shorter than intended or contained additional scaffold sequence (Figure 3b, Supplementary Figure 3). We have also tested all possible 1 nt insertions at the *AAVS1* site and observed variable ratios of intended/scaffold-containing editing products, suggesting an effect of RT template and targeted DNA sequences on PRINS outcomes (Figure 3d). Altogether, our results demonstrate that PEn can efficiently install precise insertions through NHEJ.

**Figure 3.**
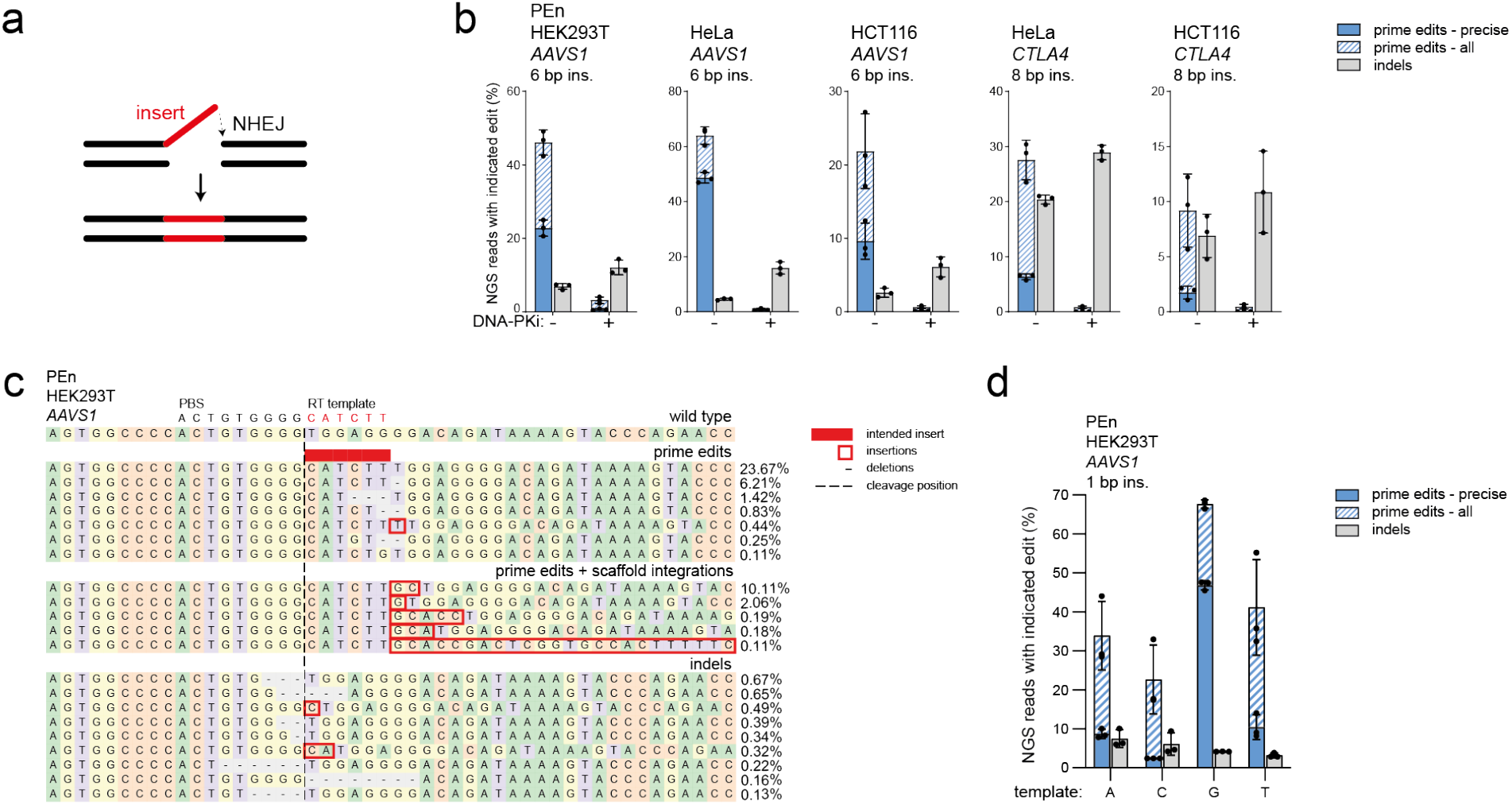
PEn editing through NHEJ. a) Model of NHEJ-mediated PEn editing using springRNA. Only DNA intermediates are shown. b) NGS analysis of PEn-mediated editing at *AAVS1* using a springRNA in HEK293T, HeLa and HCT116 cells with or without DNA-PK inhibitor. The plot shows mean ± SD of 3 independent biological replicates. “prime edits – all” and “prime edits – precise” categories are superimposed. c) Representative alignment and allele frequencies of *AAVS1* locus edited with PEn and the indicated RT template in HEK293T cells. For each category, the top 10 variants are shown with a minimum frequency of 0.1%. d) NGS analysis of PEn-mediated 1 nt insertions at *AAVS1* using springRNAs with four different RT-templates. The plot shows mean ± SD of 3 independent biological replicates. “prime edits – all” and “prime edits – precise” categories are superimposed.

### Off-target analysis of PEn editing

Integration of short double-stranded DNA fragments at DSBs has been exploited for Cas9 off-target detection ^21^ and integration of single-stranded DNA fragments was shown to increase both on- and off-target editing by Cas9 ^22^. Based on these studies, we reasoned that PEn might also show more pronounced off-target editing by actively modifying DSBs and in doing so preventing error-free DNA repair. To investigate PEn-mediated off-target editing, we have targeted three sites (*FANCF, HEK3 and HEK4*) with gRNAs that were previously profiled for off-target editing with both Cas9 and PE2 ^18,21^. We used PEn and a matching PEn mutant (PEn-dRT) carrying previously reported RT-disabling mutations ^18^ with either pegRNAs or springRNAs against the three targets in HEK293T cells and analyzed both on-target editing and a total of 11 off-target sites by deep amplicon sequencing (Figure 4). Compared to PEn-dRT, PEn induced up to 2-fold higher total on-target editing. Similarly, we observed that PEn increased off-target editing across most target sites. The increase ranged from moderate at *HEK4* off-targets (1.4 to 2.3-fold), to high at *FANCF* with off-target 1 reaching up to 13-fold increase (Figure 4). Examination of editing outcomes at these sites revealed that the increase was caused by RT-mediated insertions that constituted the majority of edits across PEn-edited sites (Supplementary Figure 4). Thus, these results show that efficient priming can increase PEn-mediated off-target editing and highlight a need for stringent peg/springRNAs and/or high fidelity Cas9 enzymes to be used with PEn.

**Figure 4.**
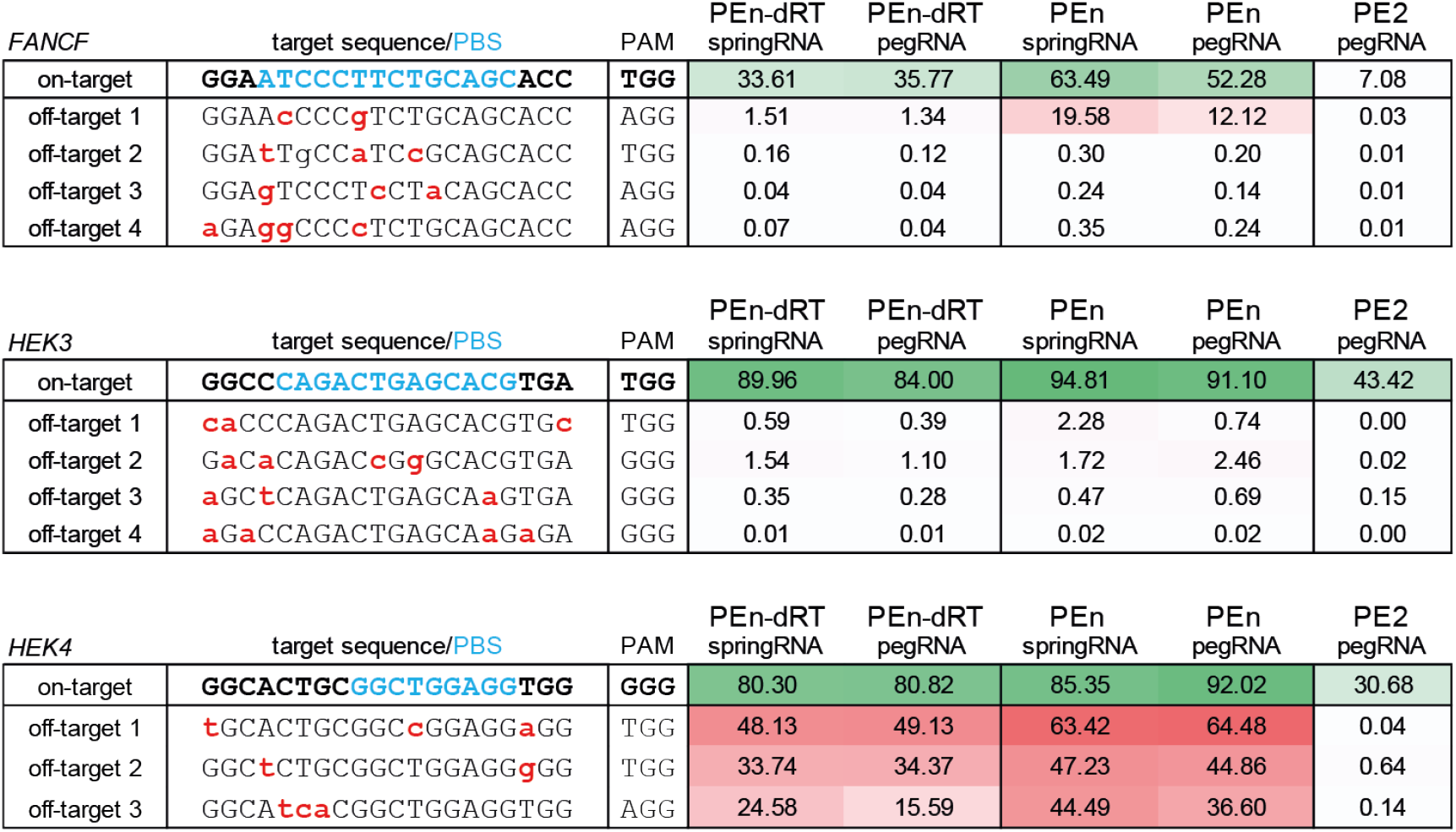
Off-target analysis of PEn editing. NGS analysis of editing outcomes at three on-target and eleven off-target sites with indicated editors and peg/springRNAs. Editing levels are shown as percentages of modified reads in each sample. The values represent average of three independent experiments. Mismatches to the on-target gRNA sequence are highlighted in red, the PBS region is highlighted in blue.

### PEn-mediated insertions at DSBs mitigate Cas9-induced large deletions

Cas9 editing has been shown to frequently cause large deletions spanning kilobase-sized regions surrounding the Cas9 target site ^23,24^. This unintended consequence of Cas9 editing poses a potential roadblock for its therapeutic applications. As PEn generates DSBs, we wondered whether PEn editing also results in similar unwanted on-target editing. To test this, we used a diphtheria toxin (DT)-based selection system in HEK293T cells ^25^ to assay for large deletions induced by Cas9, PE2 and PEn. In this system, the disruption of the *HBEGF* coding sequence generates cells resistant to DT treatment, while cells carrying an intact copy of the *HBEGF* coding sequence are efficiently killed by DT (Figure 5a). To monitor large deletions induced by different editors, we targeted an intron of *HBEGF* with either Cas9, PE2 or PEn and subjected the edited cells to DT selection. The percentage of colonies surviving DT treatment normalized to the total *HBEGF* editing levels in each condition can be used to approximate the levels of large deletions in the cell population, as only cells carrying *HBEGF* deletions larger than ~600 bp acquire DT resistance. As expected, Cas9 editing led to relatively high frequency of large deletions (Figure 5b) confirming previous observations ^23,24^. In contrast, nickase-based PE2 editing that does not induce DSBs only resulted in basal levels of large deletions. Surprisingly, similar to PE2, PEn editing with pegRNA or springRNA led to minimal levels of large deletions compared to Cas9 editing (Figure 5b), despite efficient editing at the target site (Figure S5a). We have analyzed large deletion patterns using PacBio long-read DNA sequencing ^24^ of the edited *HEBGF* locus prior to DT selection. The alignment of *HBEGF* long reads confirmed the presence of large deletions in Cas9-edited sample but not in PE2 or PEn-edited samples (Figure 5c).

**Figure 5.**
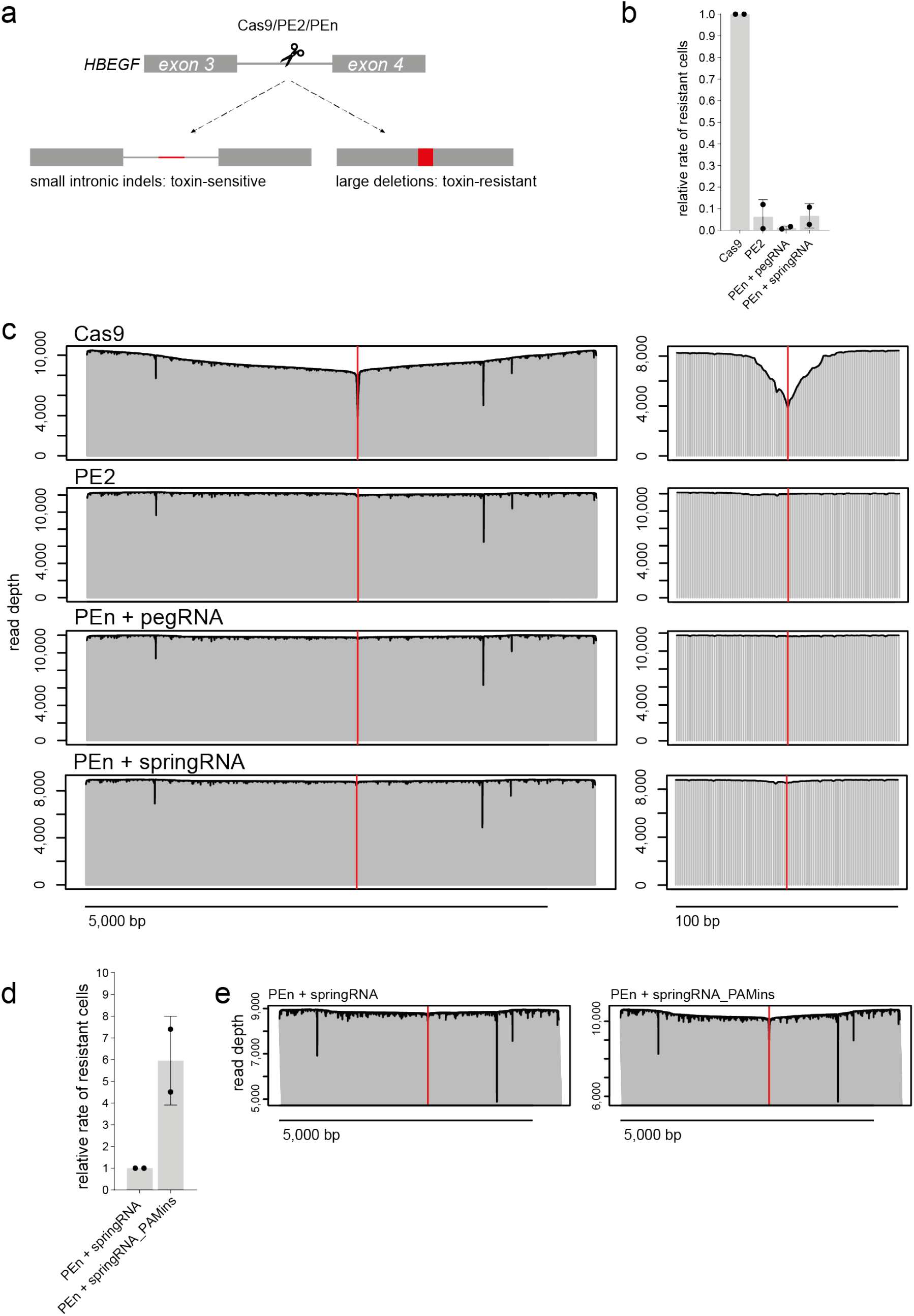
PEn-mediated insertions at DSBs mitigate Cas9-induced large deletions. a) Diphtheria toxin (DT) selection-driven assay to detect on-target large deletions induced by different genome editing systems b) Relative rates of surviving colonies after DT selection of cells edited by indicated genome editors. Data normalized to Cas9. The plot shows mean ± SD of 2 independent biological replicates. c) Alignment of long *HBEGF* reads from samples targeted with Cas9, PE2 or PEn and harvested before DT selection. Red lines denote Cas9 cleavage site. Scalebar 5000bp. Panels on the right show window of 100 bp around the cleavage site. d) Relative comparison of the rates of surviving colonies after DT selection of cells edited with PEn and either springRNA with a random insert of PAM-reconstituting springRNA. Data normalized to PEn + springRNA The plot shows mean ± SD of 2 independent biological replicates. e) Alignment of long *HBEGF* reads from samples targeted with PEn and either springRNA with a random non-PAM insert of PAM-reconstituting springRNA (PAMins) harvested before DT selection. The y-axis is set from minimal to maximal read depth for each sample.

We hypothesized that Cas9-induced large deletions might be a result of cyclic targeted DNA cutting by Cas9 after precise DSB re-ligation by NHEJ. Since PEn does not rely on random indel generation by endogenous DSB repair system, it efficiently disrupts this cycle by destroying the gRNA binding site upon successful RT-templated DNA insertion. To test this model, we have designed a springRNA encoding an insertion which reconstitutes the *HBEGF* gRNA binding site (PAMins), potentially allowing multiple rounds of PEn-mediated cutting. Indeed, we have observed ~6-fold higher rates of DT-sensitive clones after PAMins springRNA editing relative to editing with a pegRNA encoding a random non-PAM insertion (Figure 5d). Long-read sequencing of these two samples confirmed more pronounced presence of large deletions upon PAMins editing (Figure 5e). We have also performed long-read sequencing analysis post DT-selection to examine large deletion patterns in more detail (Supplementary Figure 5). PEn editing with a non-PAM-insert springRNA revealed a deletion landscape with discrete transitions in the coverage depth, suggesting that the detected large deletions originated from a small number of resistant clones, further confirming the rarity of PEn-induced large deletions. On the other hand, PAMins editing led to a complex and heterogeneous large deletion pattern resembling that of the Cas9-edited sample (Supplementary Figure 5).

Altogether, our data show that PEn editing at the *HBEGF* locus does not induce considerable levels of unwanted large on-target deletions and thus might be a safer alternative compared to Cas9 editing. Additionally, we propose that multiple cycles of Cas9 cutting facilitated by precise repair of the target locus by NHEJ might be one of the mechanisms responsible for unwanted large deletions caused by Cas9 editing.

## Discussion

In this work, we present two different strategies to introduce precise genomic insertions using a novel *Sp*Cas9 nuclease-based prime editor PEn. We showed that PEn promotes insertions through distinct DNA repair mechanisms, expanding the current nickase-based prime editing toolbox. In the first approach, we combined PEn with canonical pegRNAs to promote a homology-dependent DSB repair leading to precise insertions. Using PEn, we efficiently introduced insertions even with pegRNAs that performed poorly with PE2, suggesting that PEn can promote a more efficient DNA editing mechanism at the targeted locus. The highly efficient PEn editing also generated undesired consequences of DSB repair, such as indels, shorter prime edits and longer than intended prime edits which we found to contain additional RT-template integrations. Similar bystander editing was also observed to various extents in the PE2 editing approach ^26^. While the presence of the unintended integrations represents a downside of PEn editing, its high robustness and efficiency might be advantageous over the existing methods in situations where a seamless 3’ end of the insertion to maintain an open reading frame of the target is not necessary, such as during the correction of frameshift mutations, gene disruption by defined stop codon integration or exon-intron junction editing. To control the DNA editing outcomes of PEn, we devised a strategy to remove the unintended prime edits by inhibiting DNA-PK, a crucial mediator of NHEJ ^7^. For several genomic targets, the DNA-PK inhibitor treatment also led to a significant increase in precise editing levels. We suspect that the mechanism of PEn editing is likely to be a type of homology-dependent end joining DNA repair. While our data suggest it might be independent of Pol θ-mediated a-EJ, future studies will elucidate the detailed mechanism.

The observation of NHEJ-mediated integrations of pegRNA RT templates during PEn editing led us to the development of the springRNAs. The springRNA does not require a homology sequence in the RT template and the intended insertion is installed through precise NHEJ. This mode of PEn editing could be of particular utility because NHEJ is a preferred type on DSB repair in most human cell types and acts independently on cell cycle progression ^3,7^. NHEJ-driven precise genome editing has proved to be a valuable tool in the past but, unlike PEn, the existing approaches rely on either separately provided DNA donors or difficult-to-control indel generation ^13,14,27^.

Off-target analysis of PEn editing revealed that peg/springRNA-priming can increase the total editing levels at off-target sites to different extents. Further systematic investigation into peg/springRNA design and optimal high fidelity Cas9 utilization will be needed to fully understand and mitigate the off-target activity of PEn.

Our surprising observation that PEn does not induce large on-target deletions might provide a substantial advantage over Cas9 editing, where frequent large deletions can be of concern, especially in therapeutic applications ^23,24,28^. Moreover, our data suggest a potential mechanism by which large deletions arise during Cas9-induced DSB generation. While the precision of NHEJ is controversial ^9,29^, our data provide further evidence that NHEJ is inherently precise and possibly enables multiple cycles of target cleavage by Cas9. This “persistent” DSB may then increase the probability of faulty DNA repair leading to large deletion generation. This is in line with the observation in human embryos where long-lasting DSBs were suggested to be a potential cause of chromosomal loss or rearrangements ^30^.

In conclusion, PEn editing is an effective method for introducing small genomic insertions and expands the spectrum of DNA repair mechanisms that can support prime editing, including NHEJ, which constitutes a major pathway of DSB repair in humans.

## Methods

### DNA constructs

PE2, PEn and *Sp*Cas9 plasmids were generated by gene synthesis (GenScript). PE2 sequence including the backbone corresponds to the previously published CMV-PE2 construct (Anzalone) (Addgene #132775). To generate PEn, the H840A Cas9 mutation in the PE2 construct was reversed to the original histidine. To generate the *Sp*Cas9 construct, RT in PEn was replaced with eGFP. PEn dead-RT construct was generated by introducing previously reported mutations in RT (M3-deadRT M-MLV RT(R110S, K103L, D200N, T330P, L603W) ^18^. pegRNA constructs were generated by customizing protospacer, PBS and RT template in the target pMA-U6-pegRNA vector (GeneArt). All pegRNA sequences used in this work are listed in the Supplementary table 1. Briefly, PCR fragments encoding pegRNAs flanked by 20 bp homology sequence matching pMA (Invitrogen) target backbone were generated by template-free PCR using two partially overlapping oligonucleotides. After PCR cleanup, the fragments were assembled into the a linearized pMA backbone using HiFi DNA Assembly Master Mix (NEB) according to the manufacturer’s protocol.

### Cell culture, drug treatments and transfections

HEK293T (ATCC CRL-3216), HEK293T *POLQ−/−* (Synthego CRISPR KO pool, >90% indels), HCT116 (ATCC CCL-247) and HeLa (ATCC CCL-2) cells were cultured at 37 °C with 5% CO_2_ in Dulbecco’s modified Eagle’s medium (Invitrogen) supplemented with 10% fetal bovine serum. All cell lines were authenticated and regularly tested for mycoplasma. For gene editing experiments, cells were transfected using FuGENE HD reagent (Promega) as per manufacturer’s instructions. For 96-well plate format, cells were seeded 24 hours prior transfection at 20,000 (HEK293T) or 10,000 (HeLa, HCT116) cells per well. Cells were transfected with 110 ng of plasmid DNA per well (55 ng of pegRNA/gRNA + 55 ng of PEn/PE2/Cas9). FuGENE:DNA ratio used for all transfections was 3:1. For larger wells, cell seeding numbers and transfected DNA amounts were scaled up accordingly. Cells were harvested for gene editing analysis after 72 hours. In DNA-PKi experiments, AZD7648 (MedChemExpress, CAS No: 2230820-11-6) dissolved in DMSO was added to the growth medium 5 hours prior transfection to the final concentration of 1 μM.

### Genomic DNA extraction and sequencing analysis

Cells were harvested using Quick Extract solution (Lucigen) according to manufacturer’s instructions. Amplicons were generated using Phusion Flash High-Fidelity PCR Mastermix (F548, Thermo Scientific) in a 15 μL reaction, containing 1.5 μL of genomic DNA extract and 0.5 μM of target-specific primers with NGS adapters (primers #1-50, as listed in the Supplementary table 2). Applied PCR cycling conditions: 98 °C for 3 min, 30x (98 °C for 10 s, 60 °C for 5 s, 72 °C for 5 s). PCR products were purified using HighPrep PCR Clean-up System (MagBio Genomics). Size, purity and concentration of amplicons were determined using a fragment analyzer (Agilent). Amplicons were subjected to a second round of PCR to add unique Illumina indexes. Indexing PCR was performed using KAPA HiFi HotStart Ready Mix (Roche), 1 ng of PCR template and 0.5 μM of indexed primers in the total reaction volume of 25 μL. PCR cycling conditions: 72 °C for 3 min, 98 °C for 30 s, 10x (98 °C for 10 s, 63 °C for 30 s, 72 °C for 3 min), 72 °C for 5 min. Indexed amplicons were purified using HighPrep PCR Clean-up System (MagBio Genomics) and analyzed using a fragment analyzer (Agilent). Samples were quantified using Qubit 4 Fluorometer (Life Technologies) and subjected to sequencing using Illumina NextSeq system according to manufacturer’s instructions. For off-target analysis, amplicons were generated using Q5 Hot Start High-Fidelity 2x Master Mix (M0494, NEB). Amplicons for long-read sequencing were generated with Q5 High-Fidelity polymerase (M0492S, NEB) using primers #51-52 (Supplementary table 2) and the following PCR protocol: 98 °C 30 s 30x (98 °C 10 s 70 °C 10s 72 °C 6 min) 72 °C 6 min.

### Bioinformatic analysis

Demultiplexing of the NGS sequencing data was performed using bcl2fastq software. The fastq files were analyzed using CRISPResso2 ^31^ in the prime editing mode with the quantification window of 5 starting from the 3’ end of intended inserts. Detailed parameters are listed in the Supplementary table 3. Prime edited override sequences were used for each site. To generate the representative alignments, the window was extended to 30 to visualize homology arm integrations of different lengths. Histograms in Figure 2A were generated using CRISPResso2. Barplots were generated using GraphPad Prism 9 (GraphPad Software, Inc) or JMP 14.1.0 (SAS Institute Inc.). Long-read sequencing was performed by GeneWiz using PacBio platform. Resulting CCS reads were aligned to the reference sequence using minimap2 ^32^ (2.2.15 with “--MD -a -xsplice -C5 -O6,24 -B4” options). The resulting sam files were processed using a custom python3 script to extract read depth and location of deletions. The coverage plots were produced using R (3.4.2).

### Diphtheria toxin (DT) selection assay

To assess the rate of large deletions induced by genome editing, HEK293T cells were transfected with different combinations of PEn/PE2/Cas9 and gRNA/springRNA/pegRNA followed by a survival assay based on DT selection. In the survival assay, transfected cells (>50% confluence) were treated with DT (Sigma-Aldrich) at 20 ng/mL. Cell viability was measured using the AlamarBlue cell viability reagent (ThermoFisher) before and after DT selection. The ratio of cell viability before/after selection was calculated to indicate the rate of large deletions. Genomic DNA was harvested from each sample before and after DT selection and indel rates for each sample were analyzed by NGS.

## Acknowledgements

We thank Steve Rees, Mohammad Bohlooly and Mike Snowden for supporting this work. We thank Amelia Smith for proofreading the manuscript. This project has received funding from the European Union’s Horizon 2020 research and innovation program under the Marie Skłodowska-Curie grant agreement no. 765269 (S.W). M.P. is a PostDoc fellow of the AstraZeneca R&D PostDoc program.

## Author Contributions

M.P. and M.M. conceptualized the study. M.P., N.A., S.L. and S.W. performed most of the experimental work with help from P.H., D.D., J.B., S.v.d.P., P.M-G., S.Š and G.S. M.F. performed bioinformatical analyses. M.P. prepared the manuscript with input from all authors. M.M. supervised the study.

