## Supplementary figures for "Harnessing DSB repair to promote efficient homology-dependent and -independent prime editing"

### Supplementary figures 1-5

### Supplementary Figure 1: SpCas9 nuclease-based prime editing

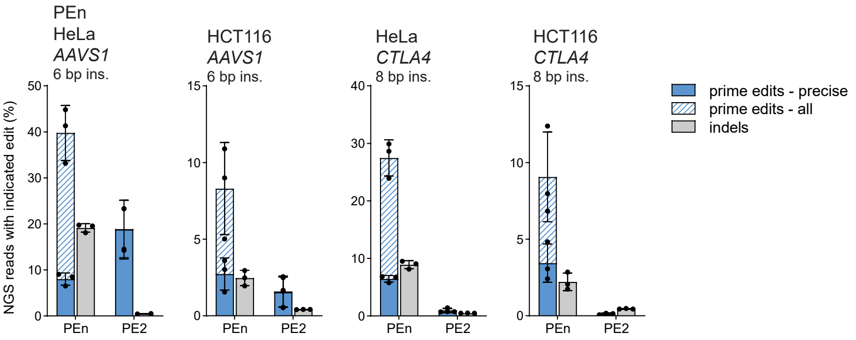


**Supplementary Figure 1** – NGS analysis of PEn or PE2-mediated targeted DNA insertions of indicated sizes in HeLa and HCT116 cells. Plots show mean ± SD of 3 independent biological replicates. “prime edits – all” and “prime edits – precise” categories are superimposed.

Supplementary Figure 2: Mechanism of PEn-based prime editing

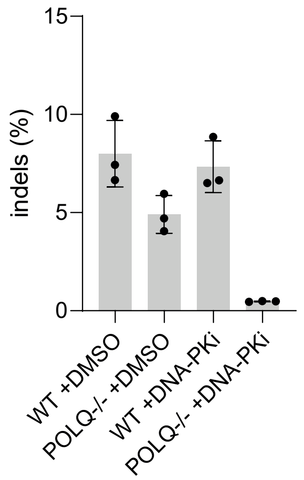


**Supplementary Figure 2** – NGS analysis of Cas9-induced indels at *AAVS1* in *POLQ-/-* cells with or without DNA-PK inhibition. Plots show mean ± SD of 3 independent biological replicates.

Supplementary Figure 3: PEn editing through NHEJ

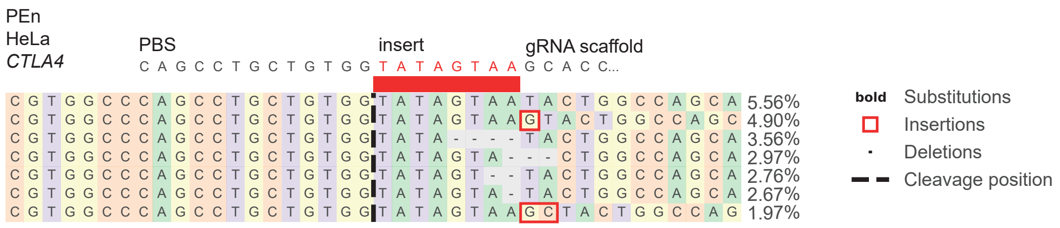


**Supplementary Figure 3** – Representative alignment and frequencies of prime edited alleles of *CTLA4* locus edited with PEn and the indicated RT template in HEK293T cells.

Supplementary Figure 4: Off-target analysis of PEn editing

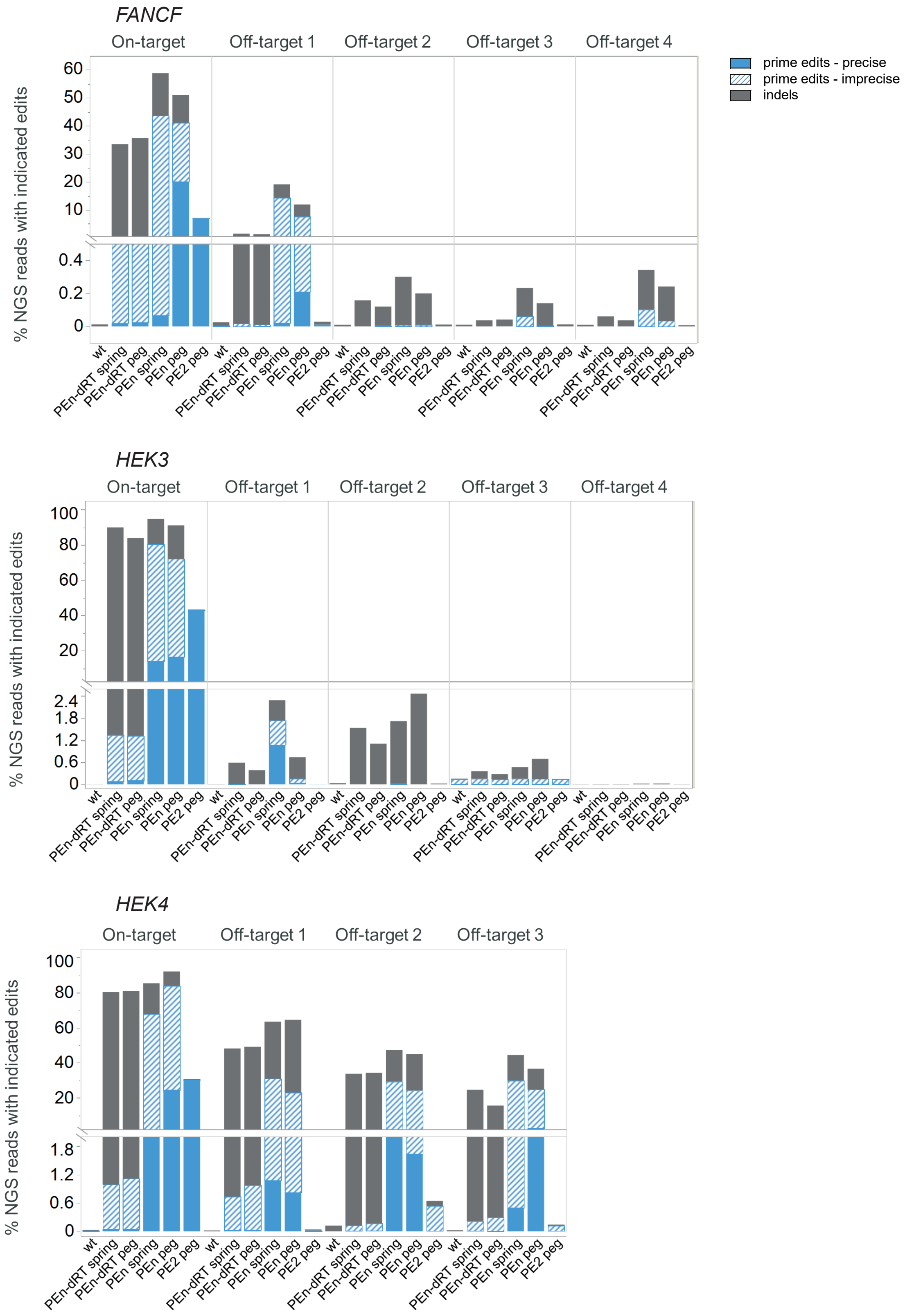


**Supplementary Figure 4** – NGS analysis of editing outcomes at three on-target and eleven off-target sites with indicated editors and peg/springRNAs. Plots show average values of 3 independent biological replicates. Indicated editing categories are stacked.

Supplementary Figure 5: Large on-target deletion induction by Cas9, PE2 or PEn editing

a

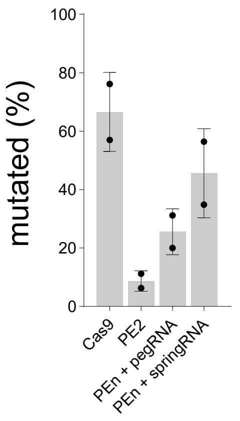


b

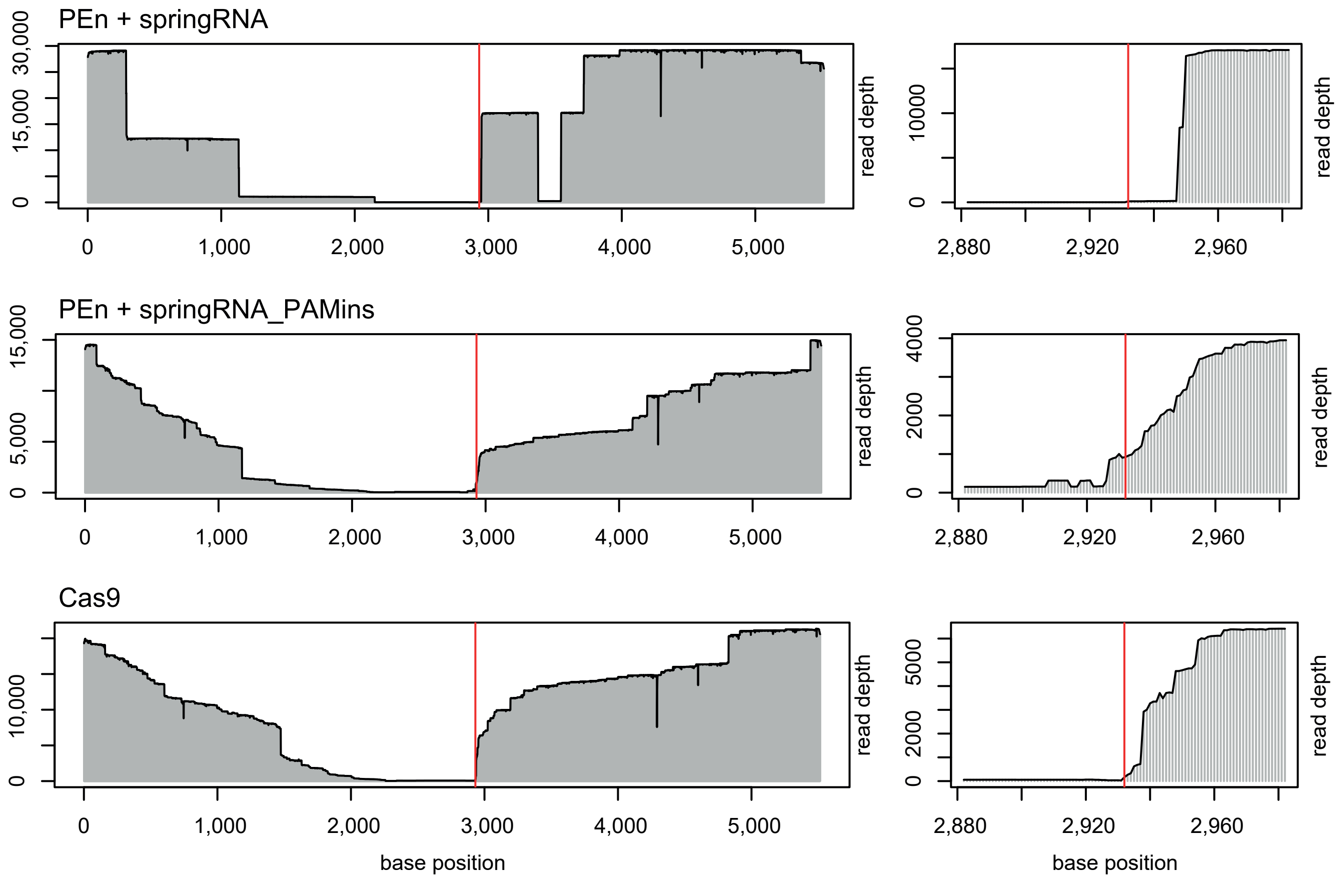


**Supplementary Figure 5** – **a)** NGS analysis of overall editing levels at the *HBEGF* locus in cells edited with Cas9, PE2 and PEn that were used for large deletion analysis. **b)** Alignment of long *HBEGF* reads from samples targeted with Cas9, PE2 or PEn and harvested after DT selection. Red lines denote Cas9 cleavage site.
